# Defining the Syrian hamster as a highly susceptible preclinical model for SARS-CoV-2 infection

**DOI:** 10.1101/2020.09.25.314070

**Authors:** Kyle Rosenke, Kimberly Meade-White, Michael Letko, Chad Clancy, Frederick Hansen, Yanan Liu, Atsushi Okumura, Tsing-Lee Tang-Huau, Rong Li, Greg Saturday, Friederike Feldmann, Dana Scott, Zhongde Wang, Vincent Munster, Michael A. Jarvis, Heinz Feldmann

## Abstract

Following emergence in late 2019, SARS-CoV-2 rapidly became pandemic and is presently responsible for millions of infections and hundreds of thousands of deaths worldwide. There is currently no approved vaccine to halt the spread of SARS-CoV-2 and only very few treatment options are available to manage COVID-19 patients. For development of preclinical countermeasures, reliable and well-characterized small animal disease models will be of paramount importance. Here we show that intranasal inoculation of SARS-CoV-2 into Syrian hamsters consistently caused moderate broncho-interstitial pneumonia, with high viral lung loads and extensive virus shedding, but animals only displayed transient mild disease. We determined the infectious dose 50 to be only five infectious particles, making the Syrian hamster a highly susceptible model for SARS-CoV-2 infection. Neither hamster age nor sex had any impact on the severity of disease or course of infection. Finally, prolonged viral persistence in interleukin 2 receptor gamma chain knockout hamsters revealed susceptibility of SARS-CoV-2 to adaptive immune control. In conclusion, the Syrian hamster is highly susceptible to SARS-CoV-2 making it a very suitable infection model for COVID-19 countermeasure development.

**One Sentence Summary:** The Syrian hamster is highly susceptible to SARS-CoV-2 making it an ideal infection model for COVID-19 countermeasure development.

## Introduction

Since emergence of SARS-CoV-2 in late 2019, the virus has spread across the globe causing >31 million confirmed infections resulting in over 960,000 deaths. SARS-CoV-2 causes coronavirus disease (COVID)-19, which is associated with a broad range of symptoms. These symptoms are most commonly fever, dry cough and fatigue, but can also include myalgia, headache, loss of taste or smell, sore throat, congestion, runny nose, nausea and diarrhea. The incubation period of SARS-CoV-2 ranges from 2-14 days with 5-6 days being most common (1). While the majority of infections are asymptomatic or present as mild to moderate cases, a small percentage of patients will progress into acute respiratory disease with fatal outcome (2, 3). In the absence of a licensed vaccine and only limited treatment options available, scientific and health care communities continue their efforts to rapidly find effective countermeasures for SARS-CoV-2 infections. Aside from new drug or vaccine development, repurposing and ‘off-label’ use of existing FDA-approved compounds is being heavily pursued, often omitting preclinical animal studies before moving directly into humans (4-7).

Preclinical animal models are integral to evaluating countermeasures for infectious diseases such as COVID-19. Non-human primate (NHP) COVID-19 models have been established, and several Old-World monkey species have been shown to be susceptible to SARS-CoV-2 infection. Infection in these animals results in a transient mild to moderate interstitial pneumonia, rather than the severe clinical outcomes (8-10). Ferrets have also been shown to be susceptible to SARS-CoV-2 infection resulting in mild disease with shedding from the upper respiratory tract. Efficient transmission has been documented suggesting the ferret may be a valuable preclinical model for transmission but not for severe disease (11, 12).

Development of small animal SARS-CoV-2 infection models was initially delayed, but recently both mouse and hamsters COVID-19 models have been described (13-17). The mouse angiotensin-converting enzyme 2 (ACE-2) receptor has only low affinity for the SARS-CoV-2 spike protein leading to poor binding and entry (18, 19). Initially, mouse susceptibility to SARS-CoV-2 infection was increased through transduction of the respiratory tract cells using adenovirus vectors expressing human ACE2, which lead to development of non-lethal pneumonia (20, 21). Several receptor transgenic mice have since been created, expressing human ACE2 under tissue-specific promoters or the endogenous mouse ACE2 promoter (14, 22-24). All of these mice are susceptible to SARS-CoV-2 infection, resulting in a range of clinical signs with mild to fatal disease depending on the transgene. SARS-CoV-2 adaptation to mice has also been attempted through either serial passaging or reverse genetics. So far this approach has sensitized mice to infection, leading to very mild disease (13).

Syrian hamsters have been reported to be susceptible to SARS-CoV-2 infection developing moderate interstitial pneumonia leading to transient mild to moderate disease (15, 17). However, these studies have used varying doses of SARS-CoV-2 for inoculation, and the impact of age and sex on infection and disease is unclear. Herein, we defined the Syrian hamster as a SARS-CoV-2 infection and disease model and refined further virologic and host parameters to increase the value of this small animal model. First, we demonstrated high functional interaction of the SARS-CoV-2 receptor binding domain (RBD) with the hamster ACE2 receptor. Next, we determined the SARS-CoV-2 dose causing infection in 50% of animals (ID_50_) following intranasal infection, showing these animals to be highly susceptible to infection. We detailed the progression of SARS-CoV-2 infection, and also the effect of sex and age on SARS-CoV-2 infection in the model. Finally, we investigated SARS-CoV-2 infection of interleukin 2 receptor, gamma chain (*IL2RG*) knockout hamsters to assess the impact of adaptive (and NK) immunity in this model.

## Materials and Methods

### Biosafety and ethics

All work using live SARS-COV-2 was performed in BSL4 using standard operating protocols approved by the Rocky Mountain Laboratories Institutional Biosafety Committee. All animal work was approved by the Institutional animal Care and use Committee and performed in strict accordance with the recommendations described in the Guide for the Care and Use of Laboratory Animals of the National Institutes of Health, the Office of Animal Welfare, the United States Department of Agriculture in an association for Assessment and Accreditation of Laboratory Animal Care-Accredited Facility. Animals were group housed in HEPA-filtered cage systems enriched with nesting material. Commercial food and water were available ad libitum.

### Virus

SARS-CoV-2 isolate nCoV-WA1-2020 (MN985325.1) was kindly provided by CDC as Vero passage 3 (25). The virus was propagated once in Vero E6 cells in DMEM (Sigma) supplemented with 2% fetal bovine serum (Gibco), 1 mM L-glutamine (Gibco), 50 U/ml penicillin and 50 μg/ml streptomycin (Gibco) (virus isolation medium). The used virus stock was 100% identical to the initial deposited Genbank sequence (MN985325.1) and no contaminants were detected.

### Cells

Vero E6 cells were maintained in DMEM (Sigma) supplemented with 10% fetal calf serum (Gibco), 1 mM L-glutamine (Gibco), 50 U/ml penicillin and 50 μg/ml streptomycin (Gibco). 293T and baby hamster kidney (BHK) were maintained in DMEM (Gibco) supplemented with fetal bovine serum, penicillin/streptomycin and L-glutamine.

### Plasmids

SARS-CoV-2 spike protein plasmids were previously described (18). Sequences from SARS-CoV-1/Urbani spike (GenBank MN908947), SARS-CoV-2 spike RBD (AY278741), human ACE2 (GenBank Q9BYF1.2) and hamster ACE2 (GenBank: XP_005074266.1) were codon optimized for humans and cloned into pcDNA3.1+.

### Cell entry assay

Vesicular stomatitis virus (VSV) particles were pseudotyped with different wildtype or chimeric spike proteins or no spike in 293T cells as previously described (18). BHK cells were transfected in 96-well format with 100 ng of host receptor plasmid, or no receptor and subsequently infected with spike-pseudotyped VSV particles as previously described (18). Approximately 18 hours later, luciferase was measured using the BrightGlo reagent (Promega), following the manufacturer’s instructions.

### Animal studies

Syrian hamsters (*Mesocricetus auratus*), 6-8-weeks-of-age and >27-weeks-of-age, males and females, were purchased from Envigo. *IL2RG* KO hamsters were generated with CRISPR/Cas9-mediated gene targeting technique established in the hamster by Utah State University as reported previously (26). Hamsters were anesthetized by inhalation of vaporized isoflurane and inoculated via intranasal instillation with 50 µl of varying concentrations of inoculum (1 to 1×10^5^ tissue culture dose 50 (TCID_50_) dropped into each naris (25ul/naris) using a pipette. Hamsters were weighed and monitored daily for signs of disease. Temperature transponders (BMDS IPTT-300) were implanted subcutaneously under anesthesia just above the shoulder blades in a subset of hamsters to monitor and record temperatures daily. Swabs (oral and rectal) were taken at different days post-infection using polyester flock tipped swabs (Puritan Medical Products). Animals were euthanized for necropsies at different timepoints to assess disease.

### Virus titration

Virus was quantified through end-point titrations performed in Vero E6 cells. Tissue was homogenized in 1ml DMEM using a TissueLyzer (Qiagen) and centrifuged to remove cellular debris (10 minutes at 8,000 rpm). Cells were inoculated with 10-fold serial dilutions of clarified tissue homogenate or whole blood samples in 100 µl DMEM (Sigma-Aldrich) supplemented with 2% fetal bovine serum, 1 mM L-glutamine, 50 U/ml penicillin and 50 µg/ml streptomycin. Cells were incubated for seven days and then scored for cytopathic effect (CPE). The TCID_50_ was calculated via the Reed-Muench formula (27).

### Viral genome load

qRT-PCR was performed on RNA extracted from blood and swabs using QiaAmp Viral RNA kit (Qiagen) according to the manufacturer’s instructions. Tissues (≤ 30 mg) were homogenized in RLT buffer and RNA was extracted using the RNeasy kit (Qiagen) according to the manufacturer protocol. Viral genomic RNA (gRNA) was detected with a one-step real-time RT-PCR assay (Quantifast, Qiagen) using primers and probes generated to target either the SARS-CoV-2 E (28) or N gene (forward: 5’-AGAATGGAGAACGCAGTGGG; reverse: 5’-TGAGAGCGGTGAACCAAGAC; probe: 5’-CGATCAAAACAACGTCGGCC synthesized with 5’ 6-carboxyfluorescein, internal Zen quencher and 3’ Iowa black quencher); all primers and probes were synthesized by Integrated DNA Technologies (IDT). Dilutions of RNA standards quantified by droplet digital PCR were run in parallel and used to calculate gRNA copies with the E assay. The N-based assay used a standard curve synthesized as follows: T7 *in vitro* transcription (ThermoFisher) of a synthetically produced N sequence (IDT) was used to generate template RNA. RNA was quantified by 260nm absorbance to determine copy number and a standard curve generated by serial dilution.

### Histopathology and immunohistochemistry

Tissues were fixed in 10% neutral buffered formalin (with two changes) for a minimum of 7 days. Tissues were placed in cassettes and processed with a Sakura VIP-6 Tissue Tek on a 12-hour automated schedule, using a graded series of ethanol, xylene, and ParaPlast Extra. Embedded tissues are sectioned at 5um and dried overnight at 42°C prior to staining. Specific anti-CoV immunoreactivity was detected using GenScript U864YFA140-4/CB2093 NP-1 at a 1:1,000 dilution. The secondary antibody is an anti-rabbit IgG polymer from Vector Laboratories ImPress VR. Tissues were then processed for immunohistochemistry using the Discovery Ultra automated processor (Ventana Medical Systems) with a ChromoMap DAB kit (Roche Tissue Diagnostics).

### Statistical analyses

Statistical analysis was performed in Prism 8 (GraphPad). T-tests were used to assess studies with 2 groups, ANOVA was used to analyze studies with >2 groups.

## Results

### The SARS-CoV-2 spike protein binds to the Syrian hamster ACE2 receptor

To validate that that SARS-CoV-2 can bind and then use as an entry receptor the Syrian hamster ACE2 receptor, a VSV pseudotype assay was performed as previously published (18). Briefly, a chimeric SARS-CoV-1 spike protein was generated with the RBD replaced with the SARS-CoV-2 RBD (VSV-SARS-CoV-2-RBD) (18). We transfected BHK cells that do not express ACE2 with expression plasmids for either human or hamster ACE2, or empty vector as a negative control. Cells were then infected with the VSV pseudotyped particles carrying either SARS-CoV-1 full-length spike (VSV-SARS-CoV-1-RBD) or chimeric VSV-SARS-CoV-2-RBD. As anticipated, both VSV-SARS-COV-1-RBD and VSV-SARS-CoV-2-RBD were unable to enter BKH cells, but entry was rescued in transduced cells expressing the hamster or human ACE2. Interestingly, VSV-SARS-CoV-2-RBD entry was increased compared to VSV-SARS-CoV-1-RBD, independent of the ACE2 origin, which may indicate higher susceptibility of both hamsters and humans to SARS-CoV-2 compared to SARS-CoV-1 (Fig. 1).

**Figure 1:**
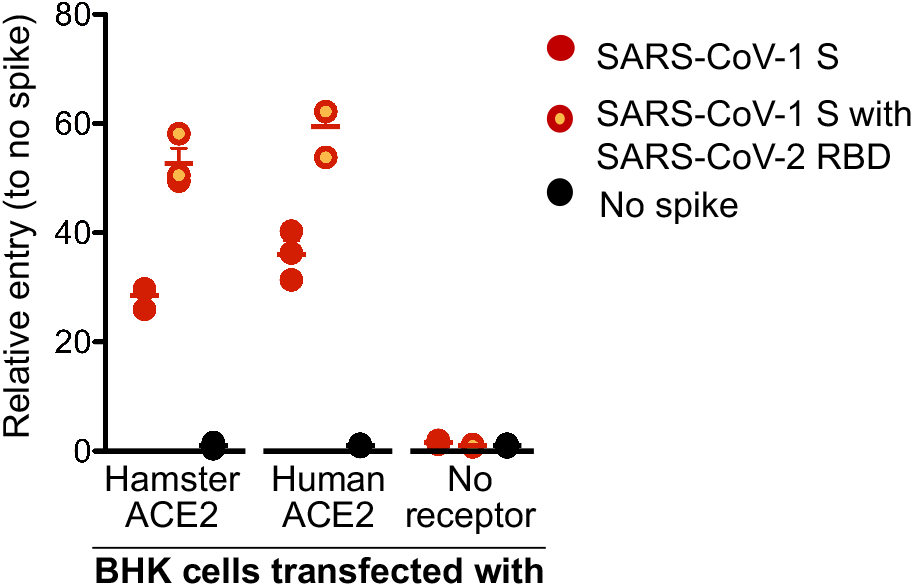
SARS-CoV-2 spike receptor binding data. A VSV pseudotype assay was used to assess the binding affinity of the SARS-CoV-2 RBD. BHK cells expressing either the human or Syrian hamster or no ACE2 receptor were infected with VSV-pseudotyped particles carrying either the SARS-CoV-1 spike (S) protein or a chimeric SARS-CoV-1 spike with the SARS-CoV-2 receptor binding domain (RBD). *Note:* red circles, SARS-CoV-1 S; yellow circles with red outline, SARS-CoV-1 S with SARS-CoV-2 RBD; black circle, no ACE-2; S, spike protein; RBD, receptor binding domain.

### Syrian hamsters are highly susceptible to SARS-CoV-2

To determine the level of susceptibility of Syrian hamsters to SARS-CoV-2 infection, four groups of six hamsters aged 4-6 weeks were infected with limiting dilutions of SARS-CoV-2 to determine the ID_50_. Groups were intranasally infected with a ten-fold serial dilution series of virus ranging from 10^3^ to 10^0^ TCID_50_, and infection course was monitored by signs of disease including weight and temperature. Although no significant changes in temperature over the experimental period were observed, weight loss between the animal groups directly correlated with the infectious dose (Fig. 2A). Oral and rectal swabs were taken at 3 days post infection (dpi) and 5dpi to measure differences in levels of viral gRNA between the different groups. Only one animal in the group receiving the 1 TCID_50_ dose had detectable gRNA in the oral swabs and none in the rectal swab (Fig. 2B), with only half the animals in this group being positive at 5dpi (Fig. 2C). Lungs were harvested at 5dpi and gRNA and infectious titers were determined. Remarkably similar gRNA levels were found across groups infected with 10 TCID_50_ or higher, only the 1 TCID_50_ group had a significant reduction in gRNA levels (Fig. 2D). Infectious titers from the lungs of each dose group had a similar pattern with high viral loads in all groups, except the 1 TCID_50_ group for which no infectious virus was detected (Fig. 2E). Remarkably, this series of experiments shows the ID_50_ in Syrian hamsters to be only 5 TCID_50_ when administered intranasally.

**Figure 2:**
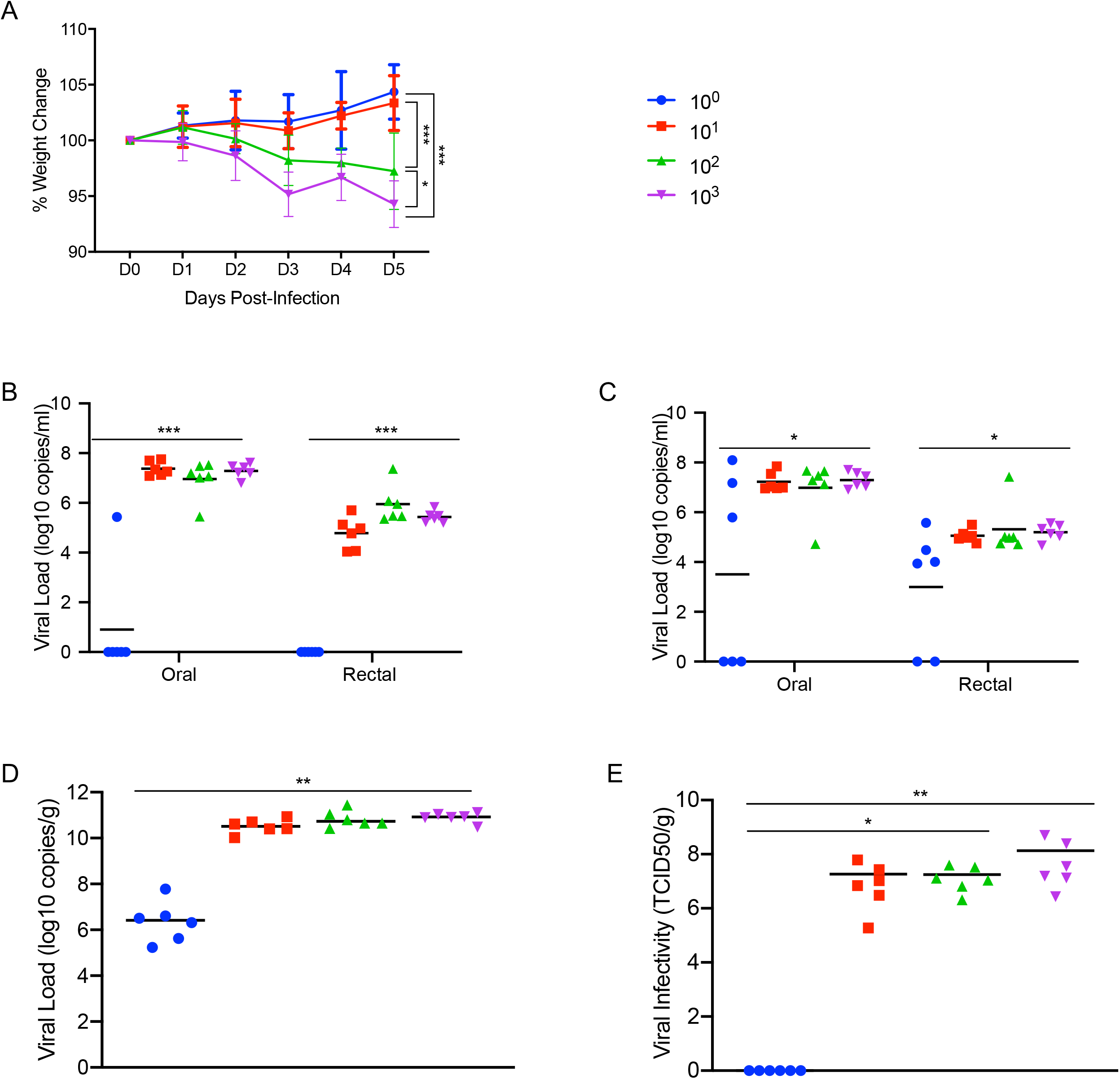
Susceptibility of Syrian hamsters to SARS-CoV-2. Syrian hamsters were inoculated intranasally with 10-fold limiting dilutions of SARS-CoV-2 beginning at 10^3^ TCID_50_. Weights were collected daily and shedding was assessed via swab samples (nasal and rectal) collected at 3dpi and 5dpi. Viral loads were determined as genome copies and infectious virus. (A) Daily weights. (B) Shedding at 3dpi. (C) Shedding at 5dpi. (D) Viral genome load in the lungs at 5dpi. (E) Infectious lung titers at 5dpi. A statistical significance was found between the groups presented in (A), with the group receiving the highest dose of 10^3^ TCID_50_ losing the most weight. The group receiving the second highest infectious dose (10^2^ TCID_50_) lost statistically less than the 10^3^ TCID_50_ group but statistically more weight than the 2 groups receiving the two lowest infectious doses. (B-E) A statistically significance difference was found between the group receiving the lowest dose (10^0^ TCID_50_) and all other groups. Multiple t tests comparing groups directly were used to analyze significance. *Note:* blue circles, 10^0^ TCID_50_ dose; red square, 10^1^ TCID_50_ dose; green triangle, 10^2^ TCID_50_ dose; purple triangle, 10^3^ TCID_50_ dose.

### SARS-CoV-2 infection of Syrian hamsters results in broncho-interstitial pneumonia

To characterize the extent of disease in this model, two groups of 10 Syrian hamsters aged 4-6 weeks were intranasally infected with either 500 ID_50_ (low dose; 10^3^ TCID_50_) or 5×10^4^ ID_50_ (high dose; 10^5^ TCID_50_) of SARS-CoV-2. Animals were monitored for clinical symptoms of disease with the intent of euthanizing a subset of animals for analysis when early symptoms became apparent. At 3dpi hamsters in both groups had lost weight on consecutive days (Fig. 3A) and displayed slightly ruffled fur with minor changes in respiration pattern. Four animals from each group were euthanized at this time (3dpi) for analysis. Lungs were examined for gross pathology and all animals had lung lesions consisting of focal extensive areas of pulmonary edema and consolidation with evidence of interstitial pneumonia characterized by a failure of the lungs to collapse following removal. No gross pathology was observed in other tissues collected including liver, spleen, kidney and brain. The remaining hamsters continued to lose weight until 5dpi with a maximum loss of <10% body weight (Fig. 3A); clinical signs remained similar until 5dpi. Three more animals from each group were euthanized at this time (5dpi). At necropsy, an increase in both number and severity of lesions were observed in animals receiving the low dose, but gross pathology was similar in appearance at 3dpi and 5dpi in animals receiving the high dose. Following four consecutive days of weight gain (Fig. 3A) and improving clinical signs, the remaining animals from each group were euthanized at 10dpi. Gross examination of the lungs at 10dpi showed a significant reduction in both lesion severity and congestion relative to lungs taken earlier during disease progression. Despite the obvious development of respiratory disease at earlier times post-infection, clinical signs were minimal at this stage (10dpi) with animals having recovered weight (Fig. 3A). Oral and rectal swabs were collected at each time point to monitor viral gRNA shedding. Shedding peaked in all swab types at 3dpi with a small decrease at 5dpi before dropping significantly at 10dpi (Fig. 3B, C). Lungs were evaluated for both SARS-CoV-2 gRNA and infectious titers. gRNA loads in the lungs were high with >10^10^ genome copies/gram (Fig. 3D), whereas gRNA loads in other organs were approximately 4-5 logs lower (Sup. Fig. 1). Lung infectious titers were 10^7^ TCID_50_ per gram at 3dpi although this had already decreased at 5dpi and was absent in all but one animal at 10dpi (Fig. 3E). Overall, there was no significant difference between the groups infected with the low and high dose of SARS-CoV-2, except for 10dpi where one animal remained positive.

**Figure 3:**
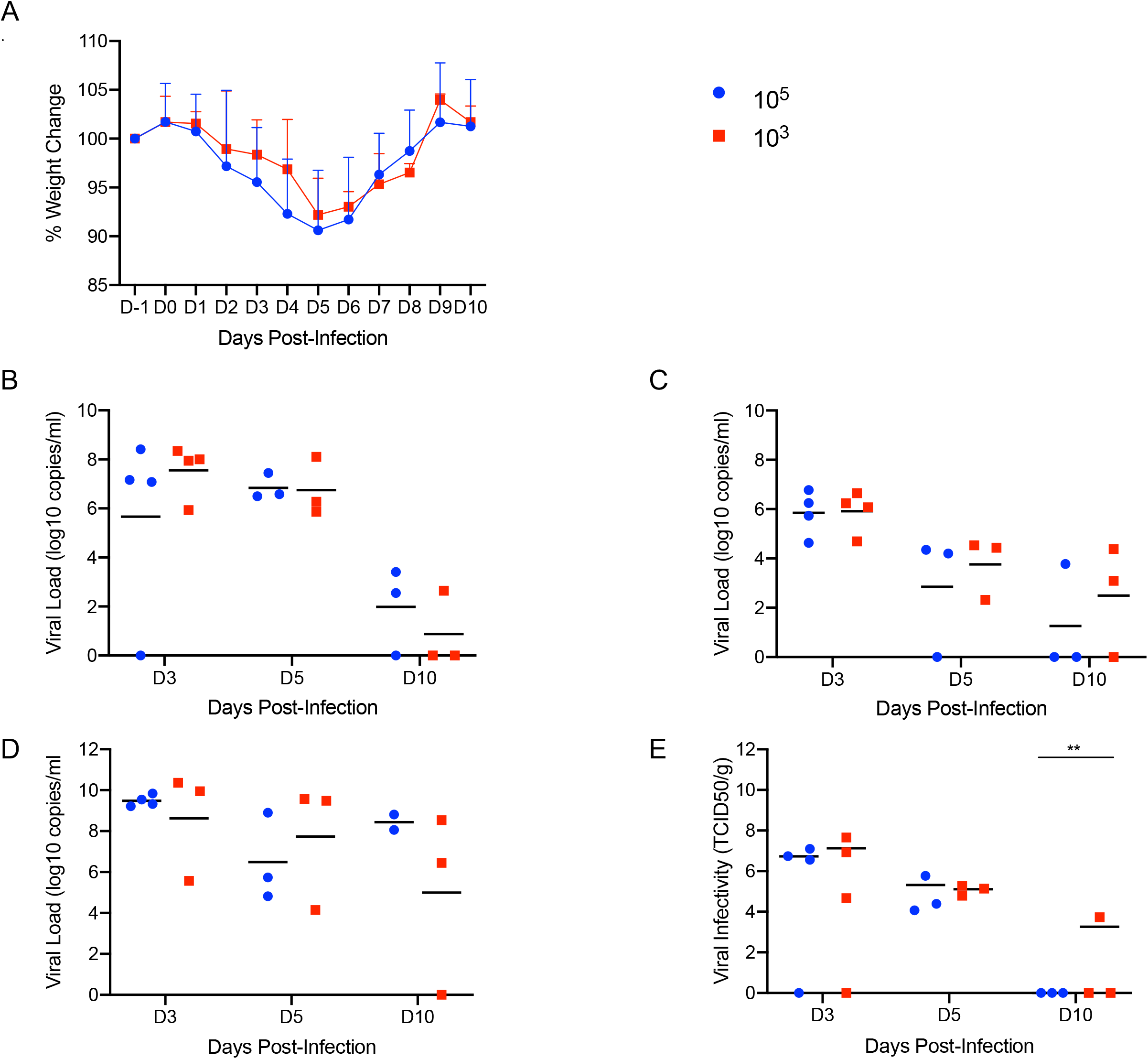
Increased infectious dose does not affect shedding or disease severity. Syrian hamsters were infected intranasally with either 500 ID_50_ (10^3^ TCID_50_) or 5×10^4^ ID_50_ (10^5^ TCID_50_) of SARS-CoV-2. Samples were collected at the time points noted. Weight were collected daily, shedding from mucosal membranes and viral genome load and infectivity in the lungs were measured. (A) Daily weights. (B) Viral genome load recovered from nasal swabs. (C) Viral genome load recovered from rectal swabs. (D) Viral genome load in the lungs. (E) Infectious titers in the lungs. T-tests were used to compare the two groups at each time where samples were collected. A significant difference was observed at 10dpi in the lung titers (E), but no other significant differences were observed in this study. *Note:* blue circles, 10^5^ TCID_50_ dose; red square, 10^3^ TCID_50_ dose.

Pathologically, changes associated with disease in the lower respiratory tract were noted in both the trachea and lung regardless of the inoculation dose. The observed pathology had a more distinct progression in the low (500 ID_50_) inoculation dose than the higher dosed group (5×10^4^ ID_50_). Evidence of broncho-interstitial pneumonia was observed at all evaluated time points. At 3dpi, lesions were characterized by epithelial necrosis in the trachea and bronchioles, squamous metaplasia of the mucosa in the trachea, bronchiolitis characterized by influx of neutrophils and macrophages into the lamina propria and mild interstitial pneumonia with expansion of alveolar septa by edema fluid, with few strands of fibrin and low numbers of leukocytes (Fig 4. A-C). By 5dpi, the interstitial pneumonia was moderate to severe with fibrin leaking into alveolar spaces, alveolar edema, influx of moderate numbers to numerous neutrophils and macrophages into alveolar spaces, presence of syncytial cells in bronchioles and alveolar spaces and prominent type II pneumocyte hyperplasia (Fig. 4D-F). Evidence of lesion resolution was observed at 10dpi with a decrease in alveolar cellular exudate, absence of epithelial necrosis and a prominent “honeycombing” pattern of type II pneumocyte hyperplasia centered on terminal bronchioles with septal expansion by a small to moderate amount of fibrosis (Fig. 4G-I). Mild and multifocal pleural fibrosis was observed at 10dpi (Fig. 5). At both 5dpi and 10dpi, moderate numbers of blood vessels were surrounded by perivascular infiltrates of lymphocytes that frequently formed distinct perivascular cuffs, occasionally focally disrupting the tunica media or forming aggregates between the tunica intima and media elevating the endothelium into the lumen. Immunohistochemical reaction demonstrated viral antigen in bronchiole epithelial cells at 3dpi (Fig. 5A-C) with fewer cells showing immunoreactivity at 5dpi (Fig. 5D-E) and no epithelial cells exhibiting immunoreactivity at 10dpi (Fig. 5G-I). Lower in the respiratory tree, SARS-CoV-2 immunoreactivity was demonstrated in type I and II pneumocytes as well as alveolar macrophages at 3dpi and 5dpi with a lack of immunoreactivity at 10dpi (Fig. 5G-I). Overall, pathologic changes progressed more rapidly in animals infected with the high dose relative to the low dose, but severity of disease was consistent between the groups with all animals developing moderate to severe broncho-interstitial pneumonia, commensurate with a mild to moderate infection model.

**Figure 4:**
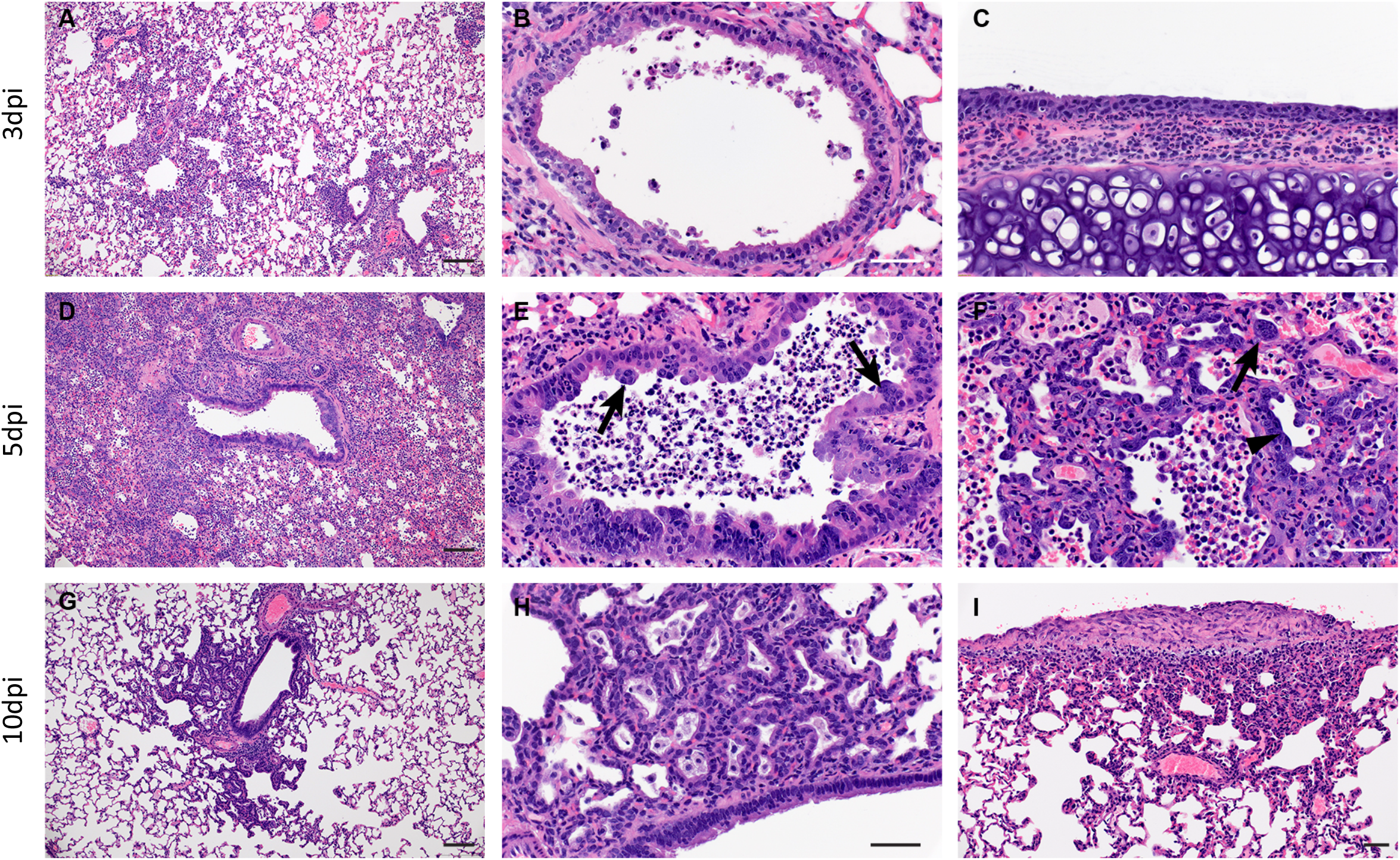
SARS-CoV-2 infection of Syrian hamsters results in broncho-interstitial pneumonia. Syrian hamsters were infected intranasally with 500 ID_50_ (10^3^ TCID_50_) of SARS-CoV-2. Lungs were fixed in 10% formalin, cut and stained with Hematoxylin and Eosin (HE) to examine pulmonary pathology at 3, 5 and 10dpi. (A-C), 3dpi. (A) Inflammation initiates within interstitial spaces in and around terminal airways with a minimal cellular exudate into the airway spaces (100x, size bar is 50um). (B) Bronchiolar epithelial necrosis with influx of neutrophils into the mucosa and airway lumen (400x, size bar is 20um). (C) Attenuation of the tracheal mucosa with loss of apical cilia accompanied with an influx of moderate numbers of degenerate and non-degenerate neutrophils (400x, size bar is 20um). (D-F), 5dpi. (D) Locally extensive inflammation is noted (100x, size bar is 50um). (E) Progressive bronchiolitis with degenerate and non-degenerate neutrophils and exudate within the lumen and prominent epithelial syncytial cells (arrows; 400x, size bar is 20um). (F) Alveolar spaces contain macrophages and neutrophils. Alveolar septa are thickened and expanded by fibrin, edema fluid and infiltrating leukocytes and are lined by prominent type II pneumocytes (arrowhead) and rare syncytial cells (arrow; 400x, size bar is 20um). (G-I), 10dpi. (G) Resolving inflammation is largely limited to bronchioles and the adjacent alveolar spaces (100x, size bar is 50um) (H) Alveolar septa are thickened by collagen with lymphocytes and lined by numerous plump type II pneumocytes that surround low numbers of foamy alveolar macrophages (400x, size bar is 20um). (I) Multifocal pleural fibrosis is evident with mild subpleural inflammation (200x, size bar is 20um).

**Figure 5:**
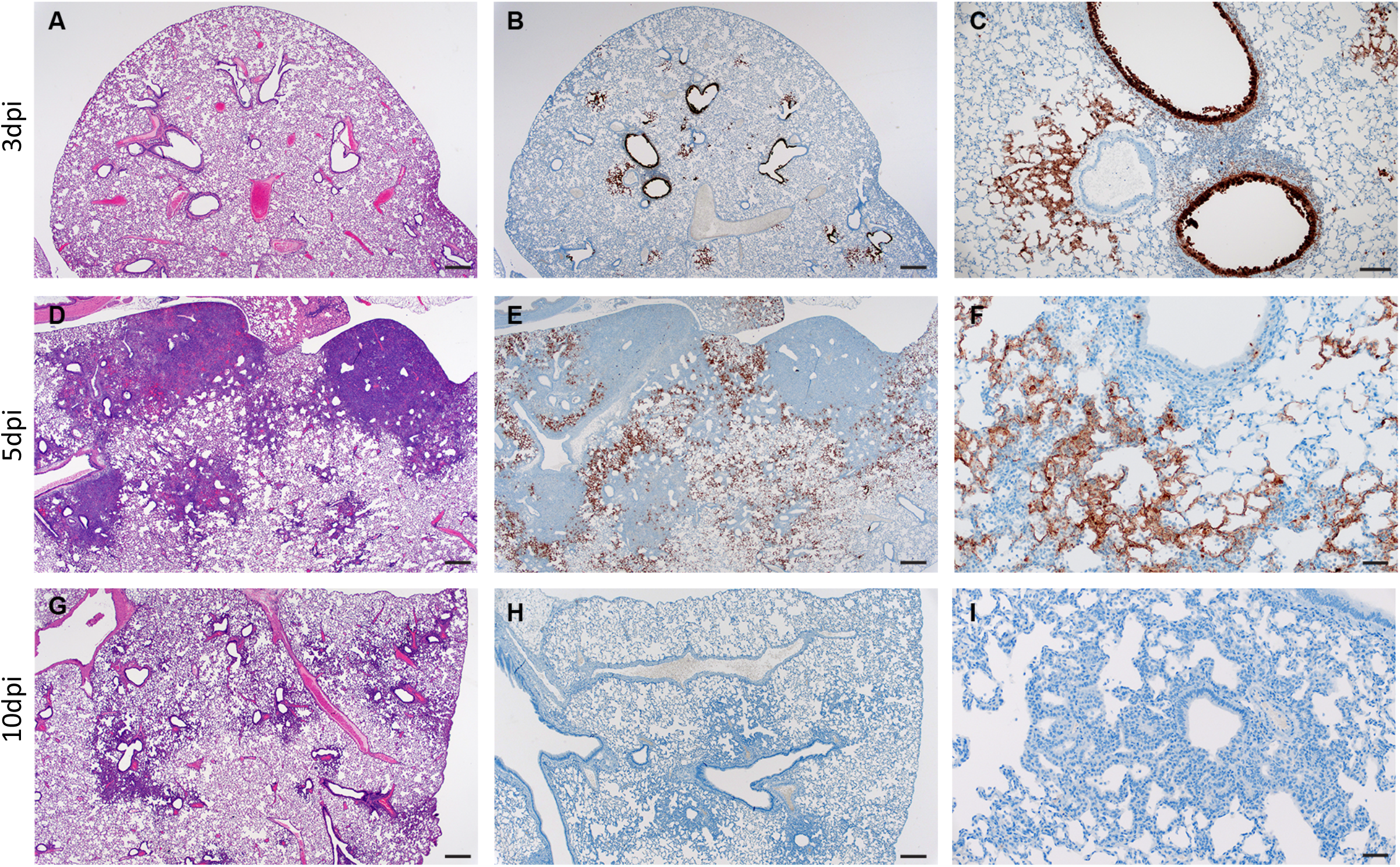
SARS-CoV-2 viral antigen in the lungs over the course of infection. Syrian hamsters were infected intranasally with 500 ID_50_ (10^3^ TCID_50_) of SARS-CoV-2. Histopathology (HE) and immunohistochemistry (IHC) was used to assess pathology with the presence of SARS-CoV-2 antigen in pulmonary sections at 3, 5 and 10dpi. (A-C), 3dpi. (A) Histopathology is largely limited to bronchioles and terminal airway spaces and is not readily apparent at a low magnification (H&E, 20x, size bar is 200um). (B) Immunohistochemical reaction highlights antigen distribution in bronchioles and terminal airway spaces (20x, size bar is 200um). (C) Bronchiolar epithelial cell immunoreactivity with limited antigen detection in alveolar spaces (100x, size bar is 50um). (D-F), 5dpi. (D) Extension of cellular exudate from bronchioles into alveolar spaces (H&E, 20x, size bar is 200um). (E) Immunoreactivity is detected along the periphery of regions of pathology and has largely been cleared from bronchiolar epithelium (20x, size bar is 200um). (F) Immunoreactivity is noted in type I and type II pneumocytes and few alveolar macrophages (200x, size bar is 20um). (G-I), 10dpi (G) Resolving inflammation is limited to bronchioles and adjacent terminal airways (H&E, 20x, size bar is 200um). (H) SARS-CoV-2 immunoreactivity is not observed in regions of resolving inflammation (20x, size bar is 200um). (I) No immunoreactivity is observed (200x, size bar is 20um).

### Neither age nor gender affected disease severity or outcome

To determine whether age or gender would affect disease progression following SARS-CoV-2 infection, we infected both young (4-6 weeks old) and aged (>6 months old) male and female hamsters with the low dose of 500 ID_50_ (10^3^ TCID_50_) by the intranasal route. Consistent with previous studies, animals began to show mild clinical signs of disease and weight loss peaking at 5dpi or 6dpi (Fig. 6A). A subset of animals in each age group and gender were euthanized for analysis at 3, 5 and 11dpi. Lung pathology was comparable in all groups at each time point throughout the study. Gross lung lesions were evident at 3dpi, had worsened by 5dpi before mostly resolving at 11dpi (Supp. Fig. 2A). To help determine disease severity, lung weights were recorded and calculated as a percentage of overall body weight for comparison. Lungs weights paralleled the observed lesions and were significantly increased at 5dpi (Supp. Fig. 2B). These observations were consistent with the development of pneumonia and was independent of sex and age.

**Figure 6:**
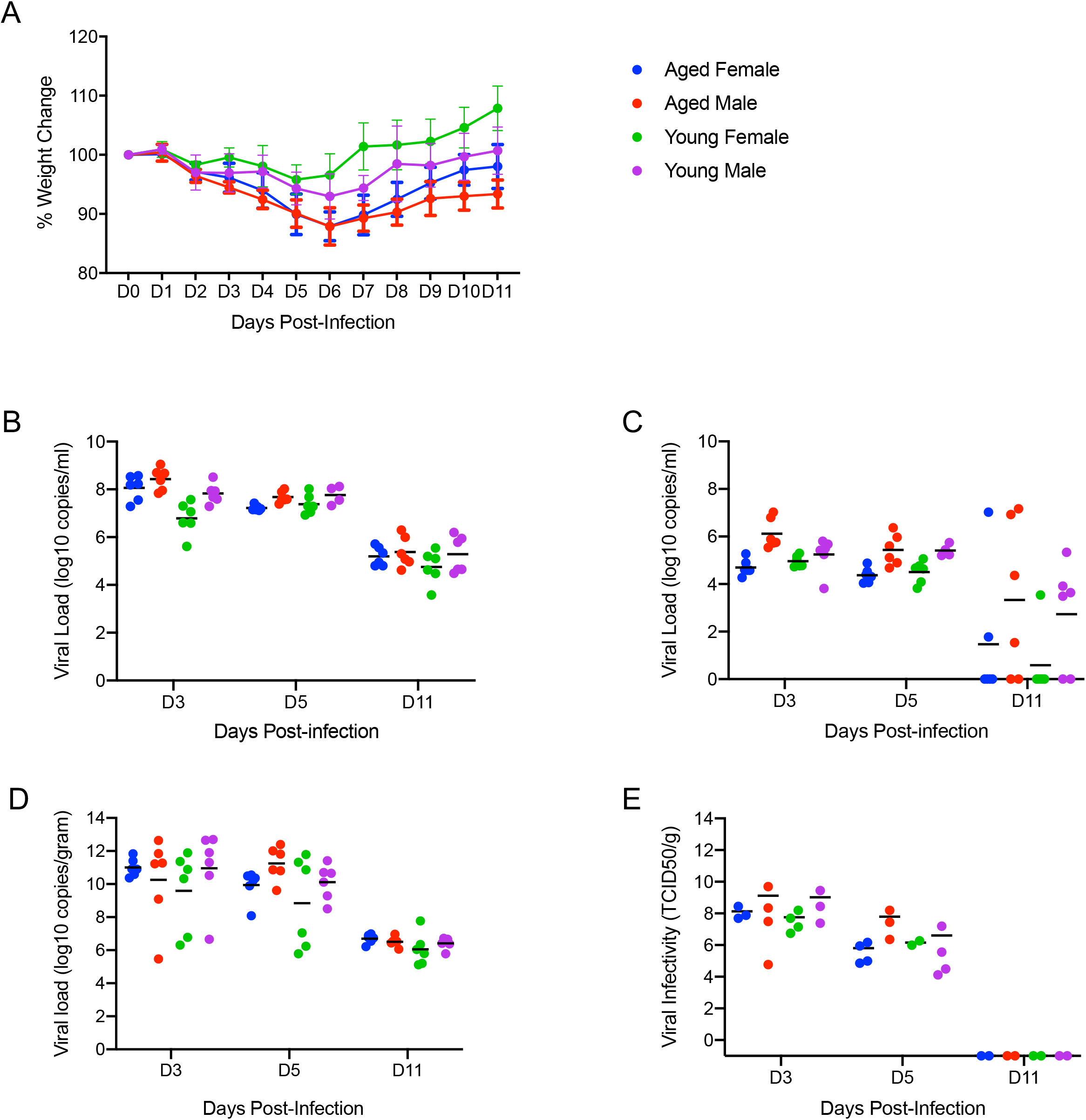
Neither age nor sex affects shedding or disease following infection with SARS-CoV-2. To compare the effects of aging and sex on disease following SARS-CoV-2 infection, young female and male (4-6weeks) and aged female and male (>6months) Syrian hamsters were infected intranasally with 500 ID_50_ (10^3^ TCID_50_) of SARS-CoV-2. Samples were collected at the time points noted. Weights were collected daily, shedding and viral loads in the lungs were measured. (A) Daily weights. (B) Viral genome load recovered from oral swabs at each time point. (C) Viral genome load recovered from rectal swabs at each time point. (D) Viral genome load recovered from lungs at each terminal point. (E) Infectious titers in the lungs. ANOVA was used to compare groups at each time where samples were collected. No significant differences were observed between groups at any time point collected in in this study. *Note:* blue circles, aged females; red circle, aged male; green circle, young female; purple circle, young male.

High levels of viral gRNA were detected in oral and rectal swabs in all groups. The highest gRNA levels detected were 3dpi before decreasing at 5dpi and again at 11dpi where only a subset remained positive (Fig. 6B, C). Interestingly, all oral swabs were positive at 11dpi suggesting viral replication was still ongoing in the upper respiratory areas (Fig. 6B). Several tissues including blood were collected and examined by qRT-PCR for viral gRNA loads at each timepoint. The lungs had high viral loads with gRNA levels highest 3dpi before decreasing at 5dpi and again at 11dpi (Fig. 6D). This corresponded with infectious titers which followed a similar pattern and peaked at 3dpi (Fig. 6E). The brain consistently had the second highest levels of viral gRNA detected and remained relatively stable across the groups at >10^8^ TCID_50_ equivalents at the time points examined. The digestive tract, both upper and lower, exhibited levels of gRNA of >10^6^ TCID_50_ equivalents across the study. Liver, spleen and kidneys all had similar levels of gRNA at 3dpi and 5dpi with >10^4^ TCID_50_ equivalents before decreasing at 11dpi.

### Hamsters lacking interleukin-2 receptor subunit gamma (*IL2RG* KO) show persistent infection with SARS-CoV-2

*IL2RG* KO hamsters are unable to develop mature NK cells with compromised development of T and B lymphocytes (26, 29). This lack of mature lymphoid cells results in an immunocompromised status known as X-linked severe combined immunodeficiency (XSCID) in humans (30). To ask the question of whether these key cellular aspects of the innate (NK) and adaptive (B and T cells) impact SARS-CoV-2 replication and associated disease, we assessed infection in *IL2RG* KO hamsters. Similar to immunocompetent Syrian hamsters, following infection with 5×10^4^ ID_50_ (high dose; 10^5^ TCID_50_) of SARS-CoV-2, four (2 males, 2 females) *IL2RG* KO hamsters lost approximately 5-10% of their body weight over the first 5 days following infection before recovering (Fig. 7A). Oral and rectal swabs were taken at 5dpi and 24dpi to measure shedding. Interestingly, both oral and rectal swabs were positive at both time-points and at very similar levels (Fig. 7B). All four hamsters were euthanized 24dpi following 2 weeks of consistent weight gain and lungs were examined for disease. At examination, the lungs had lesions similar to the immunocompetent hamsters at 5dpi, were congested and failed to collapse. Remarkably, virus titration performed on lung tissue revealed high infectious titers ranging from 10^7^ - 10^9^ TCID_50_ per gram of tissue, even at 24dpi (Fig. 7C).

**Figure 7:**
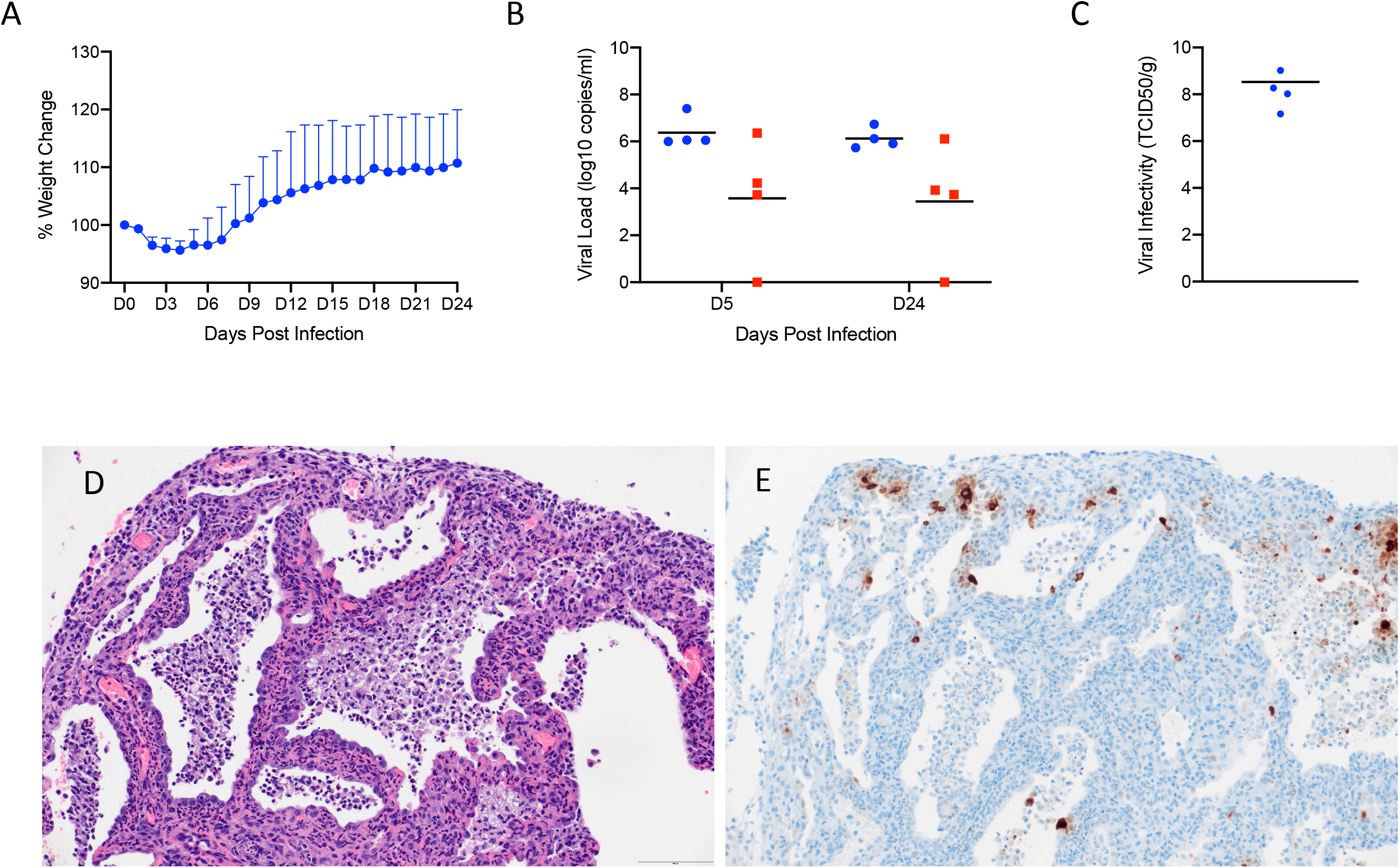
SARS-CoV-2 infection of Interleukin-2 receptor subunit gamma knockout hamsters (*IL2RG*^*-/-*^) results in persistent infection and pneumonia. *IL2RG* KO hamsters lacking mature B-cells, T-cells and NK cells, were infected with 5×10^4^ ID_50_ (10^5 TCID_50_) and followed for 24 days to determine if disease developed. Weights were collected daily and shedding from mucosal membranes and viral infectivity in the lungs were measured at the time points noted. (A) Daily weights. (B) Viral genome load recovered from oral and rectal swabs at each time point. (C) Infectious titers in the lungs. (D) Alveoli frequently contain macrophages, neutrophils and sloughed epithelial cells, and are lined by numerous hyperplastic type II pneumocytes (H&E, 200x). (E) Immunoreactivity is observed in hyperplastic type II pneumocytes and macrophages (Anti-SARS-CoV-2 nuclear protein, 200x). *Note* (B): blue circles, oral swabs; red circle, rectal swabs. Size bar is 100um.

Histopathologic analysis of the lung sections of all evaluated *IL2RG* KO hamsters exhibited disseminated, moderate to severe, chronic-active interstitial pneumonia. Alveolar septa were expanded by moderate amounts of fibrin, variably well-organized bundles of collagen and infiltrated by moderate numbers of neutrophils and macrophages. Adjacent alveolar spaces frequently contained moderate numbers to numerous macrophages with fewer degenerate and non-degenerate neutrophils admixed with cellular debris (Fig. 7D). Greater than 50% of evaluated alveolar spaces were lined by type II pneumocytes that occasionally exhibited pseudo-stratified, columnar epithelial differentiation with a distinct ciliated apical border. Lymphocyte infiltrates and perivascular lymphoid aggregates were absent in all evaluated sections.

Immunohistochemical reaction revealed numerous immunoreactive type I and type II pneumocytes as well as immunoreactive ciliated bronchiolar epithelial cells (Fig. 7E). Histopathologic evaluation of the spleen confirmed the absence of lymphoid follicles and peri-arteriolar lymphoid sheaths. Extramedullary hematopoiesis, noted in both the spleen and liver, consisted entirely of erythroid lineage cell populations.

## Discussion

SARS-CoV-2 infection in humans varies from asymptomatic to severe respiratory disease that can be fatal, especially in elderly and otherwise immunocompromised individuals. The lack of preclinical animal models that replicate the severe disease of some COVID-19 patients is a substantial hurdle for the progression of promising countermeasures from in vitro testing through to clinical trials. A reliable small animal model would allow reproducible, in-depth analyses of infection patterns, elucidation of the immune response to SARS-CoV-2 infection and serve as a critical preclinical model for the initial in vivo step in evaluating COVID-19 countermeasures for human use.

The Syrian hamster has been established as a SARS-CoV-2 animal model, but a more thorough analysis had yet to be performed. To expand our understanding the Syrian hamster model, we first determined that the hamster ACE2 receptor was compatible with the SARS-CoV-2 spike protein binding domain. The level of binding and entry was consistently higher than the corresponding assay testing human ACE2 binding activity. As the receptor binding data suggested and other recent studies have shown, Syrian hamsters were susceptible to SARS-CoV-2 infection resulting in moderate to severe broncho-interstitial pneumonia and prolonged virus shedding of at least 10 days. The ID_50_ of SARS-CoV-2 in the Syrian hamster is low, roughly five infectious particles will result in a productive infection in 50% of animals.

Symptoms of disease appear approximately 3dpi, but clinical signs are minimal with a consistent but not severe weight loss of 5-10%. Ruffled fur may be observed in some hamsters, respiration rates may increase slightly, but the behavior is unchanged from naïve hamsters. Infection was systemic following intranasal inoculation and viral gRNA was detected in all tissues examined. However, the lungs were the major site of viral replication and clearly showed a consistent but moderate pathology. Following intranasal infection, interstitial pneumonia initiating as bronchiolitis and focusing around terminal airways developed at 3dpi but was more severe in the animals receiving higher infectious doses at this early time point. Pulmonary pathology continued to increase in severity and extent of lesions in the lower dosed animals and was more severe at 5dpi, with characteristic evidence of coronaviral infection noted including presence of syncytial cells in bronchioles and alveolar spaces. Pulmonary pathology was diminished by 10dpi in all animals examined at those times with characteristic evidence of epithelial regeneration noted.

In the human population, there have been reports of an increase in COVID-19 severity in males (31, 32) as well as a disparity of COVID-19 severity among different age groups (33, 34). In these studies, neither age nor gender was a factor in disease severity or outcome. Young hamsters had similar shedding kinetics, virus titers in the lungs and developed similar pulmonary pathology as aged hamsters regardless of sex. Importantly, all animals taken at the late timepoints showed evidence of recovery from disease at a similar rate.

Interestingly, elimination of host adaptive immune responses in the *IL2RG* KO model resulted in a chronic infection persisting at least 24 days. Virus was detectable in oral and rectal swabs at 5dpi and at study termination (24dpi). Histopathology evaluated at 24dpi supports a chronic-active infection of the respiratory system with foci of epithelial regeneration as well as active recruitment of neutrophils and macrophages. Unlike the immunocompetent hamster model, in which antigen was only detectable outside of regions of pathology, SARS-CoV-2 antigen was detectable in type I and type II pneumocytes, hyperplastic pneumocytes in regions of regeneration and macrophages. Additionally, viral antigen was present in morphologically normal bronchial epithelial cells at 24dpi, a feature only routinely observed at 3dpi in the immunocompetent hamster model. While histopathology revealed a moderate to severe pulmonary inflammatory response, clinical signs of severe respiratory disease was not observed in this model. These data suggest that the innate immune system, in the background of compromised adaptive immunity, is capable of depressing the viral infection enough to keep respiratory physiology relatively stable, but incapable of eliminating SARS-CoV-2 infection.

Additionally, vascular changes and perivascular leukocyte infiltrates were not observed in the *IL2RG* KO model, unlike the immunocompetent hamster model. This data, even though limited to four animals, suggests that the adaptive immune response or IL-2 signaling pathway may play a critical role in the development of innate leukocyte recruitment and staging during viral infection and the regulation of the coagulation cascade in response to a pro-inflammatory local environment.

Both autopsy and terminal biopsy samples from human patients exhibiting COVID-19 disease have shown histologic evidence of diffuse alveolar disease, and death is frequently attributed to the clinical progression of pneumonia, often times resulting in acute respiratory distress. Diffuse alveolar disease in SARS-CoV-2 infection has characteristic lesions of hyaline membrane formation in alveolar spaces accompanied with proteinaceous fluid leaking from damaged vessels into alveolar spaces. In hamsters, there is evidence of alveolar epithelial damage at peak virus replication with lesion resolution later on. However, the hamster model fails to develop fulminant diffuse alveolar disease and lacks the respiratory decompensation associated with the clinical syndrome of acute respiratory distress. Relative to the recently developed NHP models (8, 10), the Syrian hamster model exhibits a similar mild disease phenotype and is well suited for assessing therapeutics or vaccines.

Another effective small animal model of COVID-19, the human ACE2 mouse, show these animals develop pneumonia resulting in fatal disease following SARS-CoV-2 infection (14, 24, 35). However, with this mouse model there is concern about the location and level of receptor expression in these human ACE2 transgenic mice. The increase in expression locales and levels could result in enhanced or systemic disease dissimilar to COVID-19 in humans as these animals have been reported to develop fatal encephalitis (35), a clinical manifestation not currently associated with severe COVID-19 disease in humans. Additionally, the human ACE2 mouse has been previously shown to cause neuronal death without evidence of encephalitis in the SARS-CoV-1 model (36). These complications may limit therapeutic studies in the human ACE2 mouse model.

Although, as with all animal models, there are some limitations exemplified by the lack of a systemic response to SARS-CoV-2 infection. The only mild disease manifestation and ability to quickly limit the infection make this model less suitable to study the mechanisms of severe COVID-19. However, the consistent and easily measured lung disease found in hamsters of all ages and sex make this a suitable infection model to evaluate SARS-CoV-2 countermeasure development.

## Acknowledgements

The authors thank the animal caretakers and histopathology group of the Rocky Mountain Veterinary Branch (NIAID, NIH) for their support with animal related work, and Anita Mora (NIAID, NIH) for help with the display items. This work was funded by the Intramural Research Program of the National Institutes of Allergy and Infectious Diseases (NIAID), National Institutes of Health (NIH), and partially funded through awards to The Vaccine Group Ltd, and the University of Plymouth.

## Conflict of Interest

The authors do not declare any conflict of interest.

## DISCLAIMER

The opinions, conclusions and recommendations in this report are those of the authors and do not necessarily represent the official positions of the National Institute of Allergy and Infectious Diseases (NIAID) at the National Institutes of Health (NIH).

## Figure legends

**Figure S1:**
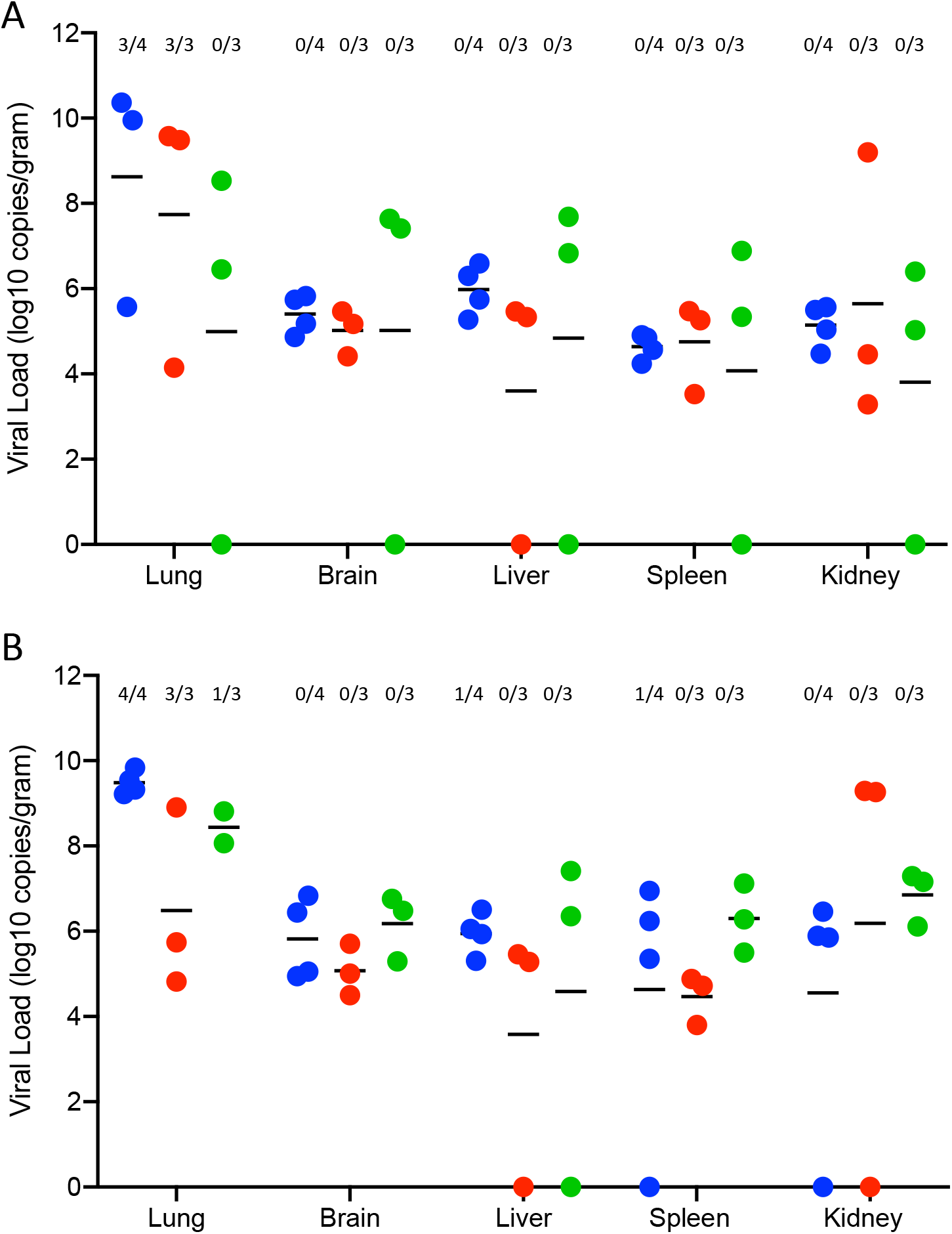
Viral genomic RNA from various tissues. Syrian hamsters were infected intranasally with either 500 ID_50_ (10^3 TCID_50_) or 5×104 ID_50_ (10^5 TCID_50_) of SARS-CoV-2. Tissue samples were collected at the time points noted. Numbers above each tissue represent the number of tissues that infectious virus was isolated from. (A) Viral genome load recovered from tissues at each time point following infection with 500 ID_50_ SARS-CoV-2. (B) Viral genome load recovered from tissues at each time point following infection with 5×10^4^ ID_50_ SARS-CoV-2. Statistical analysis using multiple T-tests found no significant differences between either group at any time point samples were collected. *Note:* blue circle, 3dpi: red circle, 5dpi; green circle, 10dpi.

**Figure S2:**
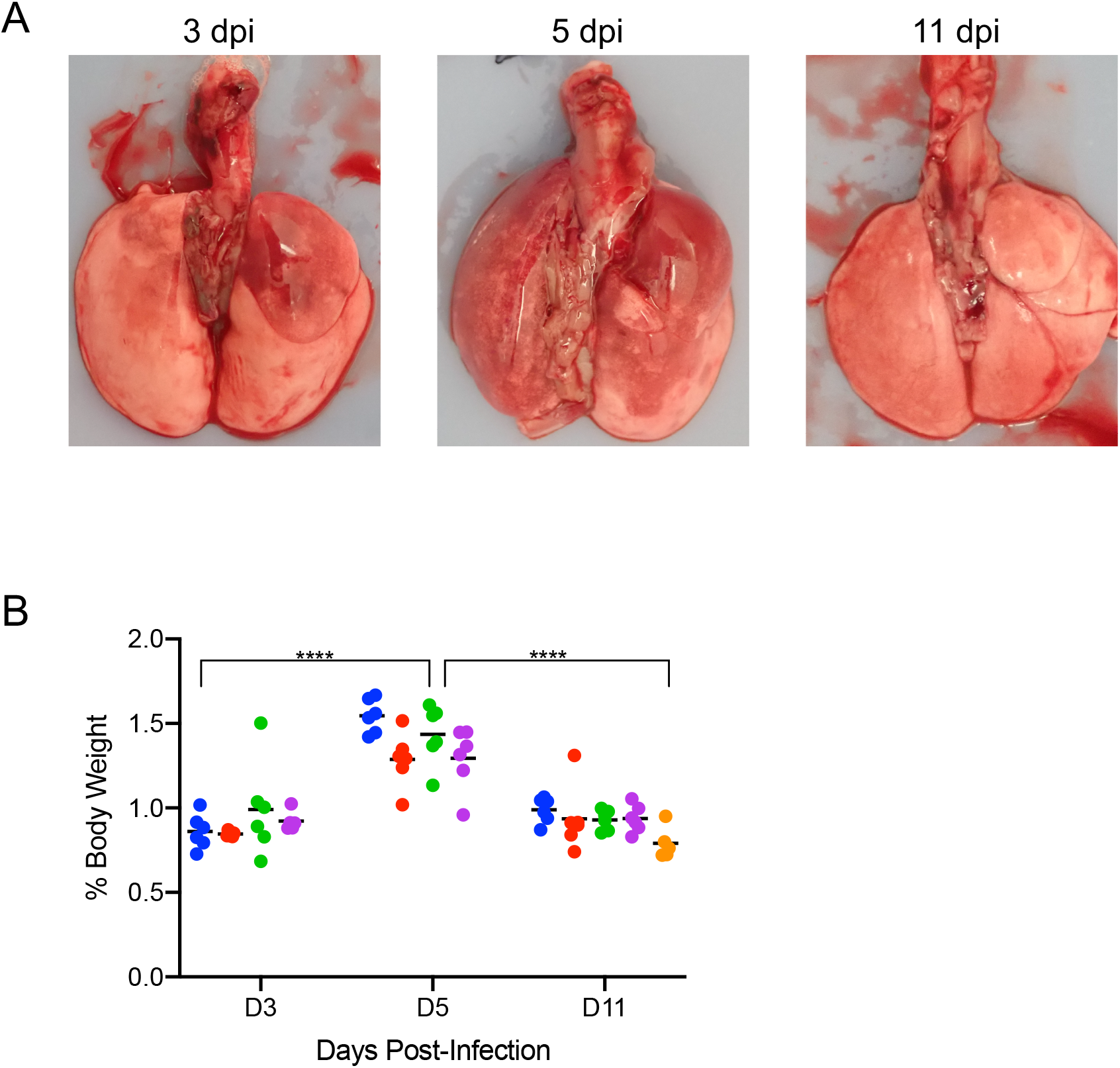
Gross pathology and lung weights. To compare the effects of aging and sex on disease following SARS-CoV-2 infection, young female and male (4-6 weeks) and aged female and male (>6months) Syrian hamsters were infected intranasally with 500 ID_50_ (10^3^ TCID_50_) of SARS-CoV-2. Samples were collected at the time points noted. (A) Representative gross pathology at indicated dpi. (B) Lung weights as percentage of body weights were recorded at each time point as a measure for pneumonia. ANOVA tests were used to compare groups at each time where samples were collected. Significant differences were not found between groups sampled on the same day. Significant differences in lung weights were found in the groups sampled at 5dpi vs those at 3dpi or 11dpi. *Note:* blue circles, aged females; red circle, aged male; green circle, young female; purple circle, young male; orange circle, naive.

## Notes

### Competing Interest Statement

The authors have declared no competing interest.

